# The thermal benefits of a mound-burrow system in a semi-desert Australian landscape: will this pebble fortress provide refuge from climate change?

**DOI:** 10.1101/2025.05.14.654155

**Authors:** Renee Claire Firman, Dustin Reid Rubenstein

## Abstract

Deserts are expected to warm and dry more than other ecoregions, making it critical to understand how arid-adapted animals might cope with increasing climate change. Plasticity in microclimatic niches created through the extended phenotype have been hypothesized to buffer desert-dwelling organisms from rapidly changing conditions. A unique example of the extended phenotype is the mound-burrow system of the western pebble mouse (*Pseudomys chapmani*), a small rodent that survives in one of Australia’s harshest environments. Here, using thermo-loggers and by obtaining mound measurements, we show that the mound-burrow system provides protection from prevailing desert conditions of both searing heat and freezing cold. We show that burrow depth and the height of the parapet around the entrance help maintain stable temperatures by warming in winter and cooling in spring. Burrows also had stable humidity levels relative to external fluctuations. We conclude that this unique pebble fortress is a critical resource that allows the species to persist in an extreme environment. We then applied modelling to assess how global warming has, and may continue to, influence(d) thermoregulation and behaviour, revealing that future summers in this extreme region may require refuge at an unachievable depth. Our study highlights the limits of the extended phenotype in buffering climate change impacts and raises concerns about the future of desert rodent populations in a rapidly warming world.

## 1. Introduction

Broadly defined as “deserts”, a large area of the earth’s surface consists of semi-arid, arid and hyper-arid land [1]. Often considered to be barren or lifeless, deserts contribute to a decent proportion of the earth’s carbon storage and flux [2, 3] and support substantial biodiversity [4]. Due to the harsh environmental conditions of high levels of solar radiation and water limitation, as well as soil salinity and nutrient deficiency, life is profoundly challenging for desert organisms. Notably, extreme temperature fluctuations are a significant abiotic stressor for desert dwelling animals [5]. Intense solar radiation, low cloud cover and minimal vegetation can result in deserts heating rapidly to daytime temperatures exceeding 40°C and dropping abruptly overnight to below freezing (<0°C) [5].

The ability to survive such extreme conditions is typically not within the scope of plastic responses, so it is generally assumed that desert-adapted species have experienced strong selection acting on complex phenotypes [5–7]. Research has demonstrated how adaptation has allowed different taxa to persist with the environmental challenges that life in the desert entails [8]. For mammals specifically, deserts present extraordinary physiological demands. Regardless of the ecoregion that they occupy, most mammals need to maintain a core body temperature of ∼36-38°C [9]. Desert dwelling mammals are thus faced with two primary thermoregulatory problems: the need to keep cool during the heat of the day and the need to stay warm overnight [10].

“Evaders” are species that avoid extreme temperatures through behaviour [8, 11]. Small desert mammals are typical evaders in the sense that they do not rely on evaporative cooling to offload excessive heat (as seen in larger mammals), instead avoiding day temperatures by only being active at night [11]. Nocturnality, however, does not necessarily equate to the complete avoidance of elevated temperatures. For example, during peak summer in the Sonoran Desert, afternoon temperatures of ∼48°C drop to an average 35°C overnight, with only one hour of the night dropping below 30°C [12]. Thus, even nocturnal “evaders” may be required to cope with elevated temperatures during the summer months [12]. Those that can only survive limited exposure are likely to adjust the duration of their activity period on a seasonal basis [12].

Burrow construction is another common behavioural strategy that desert animals use to not only evade the heat of the day, but also to survive overnight temperatures that can drop below freezing. Primarily due to differences in burrower morphology (e.g., size, scraping limb musculature, etc.), soil properties (e.g., hardness, water content, etc.) and the number of inhabitants (i.e., solitary vs. social species), the size, shape, and complexity of burrows varies considerably among species [13]. For example, the burrows of Australian desert rodents can range from simple, shallow tunnels, like those formed by the sandy inland mouse (*Pseudomys hermannsburgensis*), to complex, multi-chambered systems as constructed by spinifex hopping mice (*Notomys alexis*) [14, 15]. Burrow architecture has been shown to play a major role in thermoregulation across different taxa, and architectural features that influence buffering from surface conditions (e.g., burrow depth, burrow orientation, number of entrances, etc.) can be adjusted to create microclimates closer to optimum [16–19]. As an example of the extended phenotype, burrows represent a trait that goes beyond the individual into the surrounding environment [20]. Recent attention has been given to extended phenotypes in view of how they may buffer or amplify the effects of climate change via alterations of the microclimate that they create [21, 22]. Mammals appear to be especially ingenious when it comes to engineering a burrow in response to climatic changes [19, 23–25].

Australia’s pebble mice (family Muridae, genus *Pseudomys*), which are of a size and appearance that is very typically mouse-like (Fig. 1c), are known for the construction of unique homebases. These mice excavate a complex subterranean burrow system in rocky substrates that are crowned by a “pebble mound” at ground level (Fig. 1b; Fig. S1). Many mounds have likely persisted for tens or hundreds of years with each one consisting of thousands of pebbles, which is suggested to be the result of continutal excavation by successive generations [26]. Each of the four pebble mouse species has a discrete, non-overlapping distribution [15]. The western pebble mouse (*Pseudomys chapmani*; herein “pebble mouse”) has a distribution that covers an area of the Pilbara region of Western Australia (Fig. 1a) [27]. The Pilbara region, which represents an ancient land surface that was subaerially exposed during the Precambrian (>540 Ma), is recognised to be one of Australia’s biodiversity hotspots [28]. The Pilbara is also a hub of anthropogenic disturbance, primarily due to iron ore mining, which has intensified in recent times [28]. In addition to habitat destruction from open-cut mining practices, Pilbara fauna is threatened, both directly (e.g., predated upon, competition over resources, etc.) and indirectly (e.g., soil compaction, exposure to predators, etc), by feral species [28]. On top of this, climate change is expected to be an especially serious threat to organisms in the Pilbara [28].

**Figure 1.**
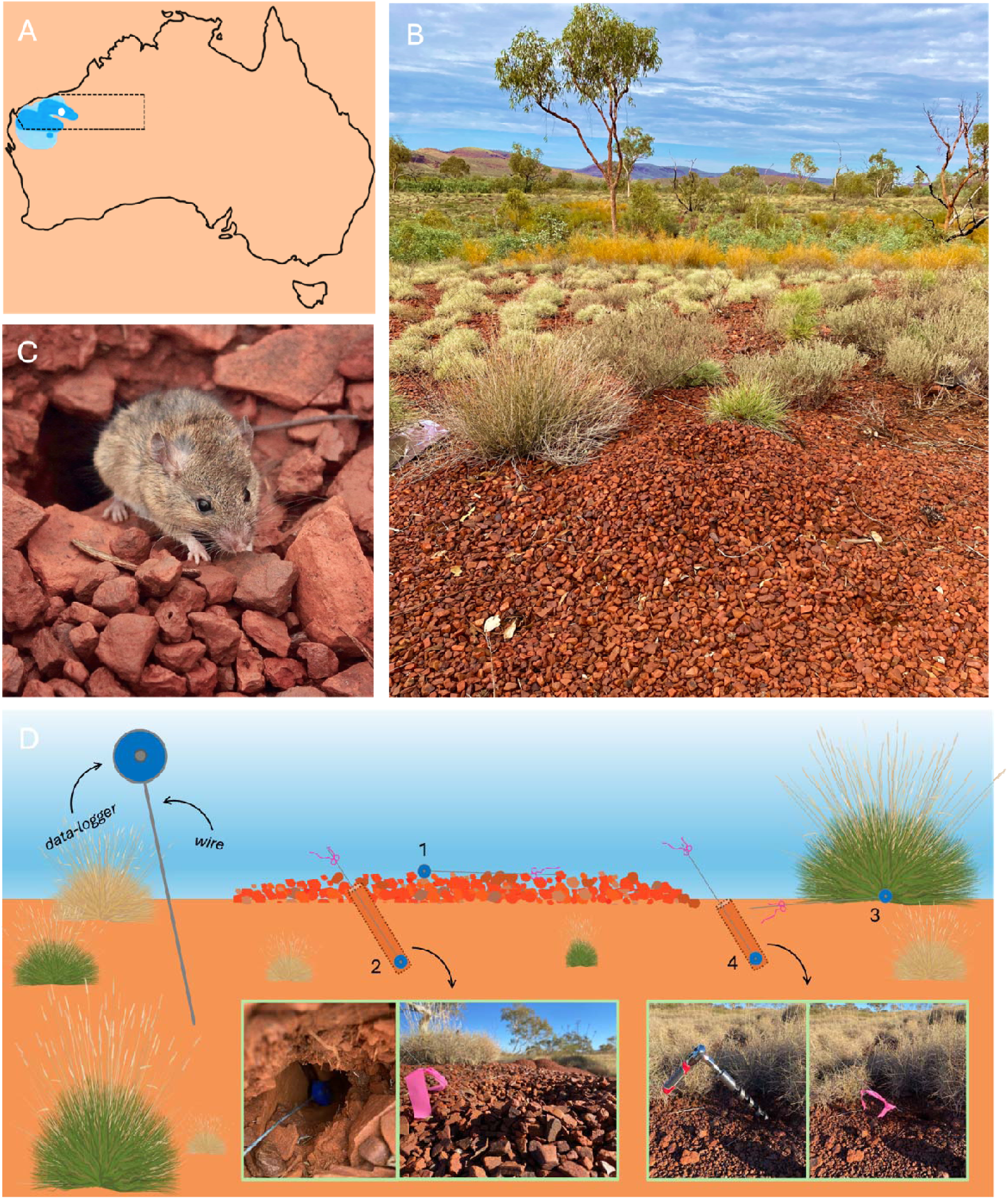
Details of the investigation, including the approximate location of the study site indicated by the white dot within the Pilbara region (dashed lines) of Western Australia; the historical distribution (light blue shading) and current distribution (dark blue shading) are shown (A). An example of a pebble mound displaying prominent features including parapets that surround burrows (photo: an) (B). A western pebble mouse at the edge of a burrow after being released from a trap (photo: (C). An overview of the investigation design displaying the locations of the thermo-loggers, including a ground logger (1) and three shelter loggers: mound burrow (2), spinifex shade (3) and experimental burrow (4; winter only) (D).

The Pilbara experiences some of Australia’s most extreme climatic conditions. Most days of the year are hot, sunny and dry, and rainfall that predominantly falls during the summer months is unpredictable and highly variable. With maximum daily summer temperatures often >43°C, the region has recorded Australia’s highest annual temperature in almost half the years over the past two decades, and temperatures are set to rise [28, 29]. Compared to a 1961-2012 baseline period, climate change models project that Pilbara temperatures will increase 1.8°C by 2030 and 2.9°C by 2050 [28, 29]. Higher temperatures means higher levels of evaporation, and this, combined with a projected 2% reduction in median rainfall by 2050 (compared to a 1961-2012 baseline period; [28, 29]) will impact the region’s fauna. Increased heat loads may lead to changes in behaviour, for example with shifts in activity patterns or a requirement to seek thermally buffered microclimates deep(er) underground [10]. These changes may allow individuals to remain within the prescriptive zone (i.e., the ambient temperature range over which body core temperature is held relatively constant), but may also result in detrimental fitness consequences like a reduction in time spent foraging or an increase in competition and/or predation [10].

Here, we combined external sensor monitoring, finescale mound measurements, and niche modelling to understand how the unique pebble mouse homebases, the mound-burrow system, provides a buffering microenvironment from some of Australia’s most extreme climatic conditions. First, we used small loggers to record burrow temperatures and humidities during logging sessions in winter and spring. Next, we measured different mound-burrow characteristics to determine the thermo-buffering properties of this component of the pebble mouse’s extended phenotype, and used camera traps to compare mouse activity patterns at these different times of the year. Finally, we used niche modelling to determine (i) how current burrow temperatures compare to historical soil temperatures at similar depths, and (ii) assess how past, present, and future conditions have, are, and will likely influence pebble mouse thermoregulation. Combining all of this information, we consider how a warming climate has the potential to lead to a narrowed prescriptive zone and changes in behaviour and activity patterns. Our findings are relevant to other arid dwelling animals across the globe, especially small mammals that are threatened by multiple stressors.

## 2. Methods

### (a) Study location and study species

The study was conducted in Karijini National Park (KNP) (22°29′46″S, 118°23′50″E/ 22.49611°S, 118.39722°E), located in the Pilbara region of Western Australia (Fig. 1a). KNP is 6,274 km² in area and is characterised by a rugged terrain of hills (700 to 800m above sea level) that form part of the Hamersley Range, as well as with deep gorges that are permanently filled with water. The climate is tropical semi-arid, with air temperatures in summer ranging from 18 to 48°C and in winter from 4 to 37°C [30]. Characteristic of the Pilbara region is highly variable annual rainfall that mainly falls in the summer (range last five years = 180–450mm; [30]). Heavy summer rainfall is often associated with cyclones that are accompanied by temperatures >40°C, with the highest temperatures occurring in January and the lowest in June. The vegetation cover is mainly composed of spinifex (*Triodia spp.*), as well has a sparse overstorey of eucalypts and shrubs, typically *Acacia*, *Senna* and *Ptilotus*.

The western pebble mouse is endemic to Australia’s Pilbara region [27] and considered to be “social” [31]. Although at lower frequencies the mound-burrow system may be occupied by multiple individuals of both sex and different ages [32], we have found that most mound-burrow systems are occupied by a single individual (typically a female; pers. obs.). The species displays reversed sexual dimorphism, with adult males being on average 30% smaller than adult females (max. weight: males = 12.5g; females = 18g) [33]. Breeding occurs over from late autumn (late May) through to the beginning of spring (early September) [26]. Pebble mice are thought to be obligate burrowers, though recent observations suggest that it may only be females that dig burrows and excavate pebbles (R, pers. obs.). An anecdotal report suggests that the pebble mouse mound-burrow system has multiple tunnels and chambers and can extend up to 60cm below the surface (electronic supplementary material Fig. S1; [34]). In addition to having to cope with severe and unpredictable environmental conditions, pebble mice have several native predators, including the Stimson’s python (*Antaresia stimsoni*) and the Australian owlet-nightjar (*Aegotheles cristatus*), as well as the introduced cat (*Felis catus*).

### (b) Mound-burrow system data collection

Mound-burrow systems with at least one open burrow (range of open burrows per system = 1–3) were selected and data-logged in winter (June 2023; n = 24) and spring (October 2023; n = 24). Because pebble mounds are typically oval in shape, we measured a long (major axis) and short (minor axis) length and calculated the area using an online ellipse area calculator (www.omnicalculator.com/math/ellipse-area). Average pebble weight was calculated by weighing 30 pebbles. One burrow per mound-burrow system was logged for temperature and humidity (30-minute intervals; 24 hours). This was achieved by following similar methodology of Moore *et al.* [21]. Temperature (°C) and relative humidity (%) data-loggers (Thermochron iButton DS1923) were taped to a ∼50cm length of high tensile wire and placed down the burrow entrance (N.B. the logger was taped in such a way that the sensor was exposed). These burrows were measured for (i) parapet height (i.e., height from the burrow entrance to the top of the parapet), (ii) aspect (i.e., orientation of burrow in relation to North), and (iii) angle (i.e., determined by using the wire extending from the burrow, a small spirit level and a protractor). We also measured the length of wire that extended into the burrow and used this to calculate burrow depth (i.e., by also using the angle of the burrow angle and the right-angle triangle formula: www.cleavebooks.co.uk/scol/calrtri.htm). A second data-logger (also taped to wire) was placed on the ground in approximately the centre of the mound (winter = 24; spring = 24). A third data-logger (also taped to wire) was placed under a spinifex bush that was close to the mound (winter = 8; spring = 24). To assess how the pebble mound may contribute to the thermal propoperties of the underground burrow system, during the winter logging session a subset of the third loggers were placed down experimental burrows (n = 16) that had been excavated to the side of the mound using an auger drill bit attached to rachet driver (Fig. 1D). The orientation, angle and depth of these experimental burrows mimicked that of the corresponding mound burrow, but lacked the pebble mouse fortress above. It was not possible to create experimental burrows in spring due to the dry, powdery nature of the earth (i.e., the tunnel collapsed when the auger drill bit was removed). Four mounds were logged during the same 24-hour period (total n_loggers_ = 12), resulting in six rotations required to acquire data for 24 mounds. Data-loggers were randomly allocated to the different positions (i.e., mound burrow, ground, experimental burrow/spinifex bush) between each rotation. Temperature and relative humidity data were downloaded from the loggers and organised for analysis. Methodology and results associated with the humidity data are provided in the electronic supplementary material.

### (c) Camera trapping

Activity data was extracted from camera traps set on mounds in the study area (n = 11). These mounds were far enough apart to allow for spatial independence according to reported home range sizes [31]. Each camera was fixed to a 130cm steel stake at a height of approximately 70-90cm from ground that was positioned at the edge of the mound on a tilted angle (Fig. S2). Dummy triggers were conducted to ensure that the camera was positioned appropriately with the entire mound surface displayed in the images. We extracted activity data for four weeks that overlapped the thermo-logging periods: (i) 7^th^ to 28^th^ June (winter) and (ii) 7^th^ to 28^th^ October (spring). As pebble mice are a strictly nocturnal species [26], the cameras were set to capture images between the hours of 17:00 to 07:00 with a burst of five images per trigger. The images were loaded into Colorado Parks and Wildlife (CPW) Photo Warehouse for pebble mouse identification [35, 36]. The CPW Photo Warehouse activity analysis was then applied to extract activity data, which included “location” (mound), “image date” and “time” (decimal time) of each image that contained a pebble mouse [35, 36]. Since we were interested in total mound activity (i.e., irrespective of the activity of discreet individuals), we did not apply a “quiet period” [36]. We also recorded detections of Stimpson’s pythons during the spring camera trapping period.

### (d) Soil temperature and thermoregulation modelling

We used the NicheMapR Biological Forecasting and Hindcasting Tools to gain an understanding of how global warming conditions may have influenced, and may continue to influence, the thermoregulation and activity patterns of pebble mice. Specifically, we used the Global Soil Microclimate Calculator to generate location specific (i.e., using the mound latitude and longitude coordinates) soil temperatures at the depths that we had logged mound burrow temperatures (http://bioforecasts.science.unimelb.edu.au/app_direct/soil/). This dataset was derived from historical data (1960-1990) that represents a typical day in each month [37]. We applied the default settings, except for adjusting the aspect to match that of each burrow and increasing the density of the soil to reflect the iron ore rich properties of the Pilbara region (mineral density: 3.6; bulk density: 1.6). We compared the historical data with our 2023 thermo-logged data. For these comparisons, we reduced the thermo-logged data to hourly temperature logs to match the hourly temperature measurements provided by NicheMapR. Further, NicheMapR provides soil temperatures at depths of set increments. Therefore, for each mound we extracted the past temperatures at the depth that most closely matched the depth at which each burrow temperature was recorded.

We used the Endotherm Model to gauge how changing conditions may have, and may continue to, influence western pebble mouse thermoregulation in relation to residing at a subterranean depth that ensures survival (http://bioforecasts.science.unimelb.edu.au/app_direct/endotherm/). For these models, we used the coordinates of a mound-burrow system that occurred at approximately the midpoint of our study site (-22.61989°, 118.35638°). We applied settings based on information that was either (i) inferred (BMR equation: mammal; shape: ellipsoid; orient to the sun: perpendicular; breath humidity: 60%; all pant parameters: 0), (ii) from trapping data (weight: 0.018kg and 0.0125kg; height: 4cm), (iii) available on thermoregulatory parameters of the arid-dwelling, and closely related, sandy inland mouse (target *T*_B_: 36°C; min. metabolic rate: 0.2W; max. *T*_B_: 42°C; min. *T*_B_: 35°C) [38], or (iv) available on *Mus musculus* (pelt depth: 5mm; max. pelt depth: 7mm; hair diameter: 0.04mm; hair length: 7mm; pelt density: 40I/mm^2^). All other parameters were set as the default; sensitivity analyses (not shown) revealed that small changes in these parameters did not change the thermoregulation profile. We obtained thermoregulation data for June (winter) and October (spring), as well as January as it is the hottest month of the year. We ran simulations for past conditions (historical dataset: 1960-1990) and applied warming conditions to model (i) current conditions as historical conditions plus 1.5°C based on a report that “Australia, on average, has warmed by 1.51±0.23°C since national records began in 1910, with most warming occurring since 1950” [39], and (ii) future conditions as historical conditions plus 3.0°C based on the expected increase in the Pilbara by 2050 [28, 29]. We ran these models for ground level and at 10, 20, 50 and 100cm underground. We also ran thermoregulation models at ground level with warming conditions of +4, +5 and +6°C for January, June and October (2100 High Emission Scenario; [40]).

### (e) Data analyses

We used descriptive statistics (mean ± s.e. or mode; minimum and maximum values) to summarise the mound characteristics data. To explore any consistent pattern in burrow aspect (e.g., an avoidance of a northern orientation), we partitioned the data into 60° categories and applied a chi-square test whether the observed frequencies differed from expected frequencies. We used linear mixed models (LMMs) in all other statistical analyses. We tested whether the magnitude of temperature buffering differed among the different shelter types (mound burrow, experimental burrow, spinifex shade). We also assessed which mound characteristics accounted for variation in mound-burrow temperatures and humidity. These analyses were performed on the entire dataset, which included “all hours” (i.e., 00:00 to 23:00), as well as in data subsets that included only the “cold hours” (i.e., 20:00 to 05:00) or the “hot hours” (i.e., 10:00 to 15:00). As there was a significant positive correlation between mean ground temperature and mean burrow temperature (electronic supplementary material Fig. S3), ground temperature was included in the LMMs testing burrow temperatures. Finally, we tested for differences between burrow temperatures and historical soil temperatures at similar depths (generated by NicheMapR). Mound ID, time point and/or rotation of recording (1 to 6) were included as random factors when appropriate. All analyses were performed in R version 4.2.0 [41].

## 3. Results

### (a) Temperature and activity

The logged ground temperatures in winter dropped below freezing overnight and reached to 35–40°Cs in the middle of the day (Fig. 2A). Notably, burrows mostly maintained temperatures of between 10°C and 20°C over the 24-hour period (Fig. 2A). As expected, temperatures in spring were higher than in winter, with ground temperatures between 15–25°C overnight and reaching up to 60–70°C in the peak of the day (Fig. 3A). The pattern of heat gain in spring was similar to what was observed in winter, though to an elevated degree with burrow temperatures not often dropping below 30°C overnight and reaching as high as 50°C in the middle of the day (Fig. 3A). In both winter and spring, the burrow temperatures took longer to cool down relative to the ground temperatures (Fig. 2A; Fig. 3A). This was especially evident in spring when ground temperatures would drop abruptly after sundown, but with the burrows cooling less rapidly and retaining most of the heat overnight (Fig. 3A).

**Figure 2.**
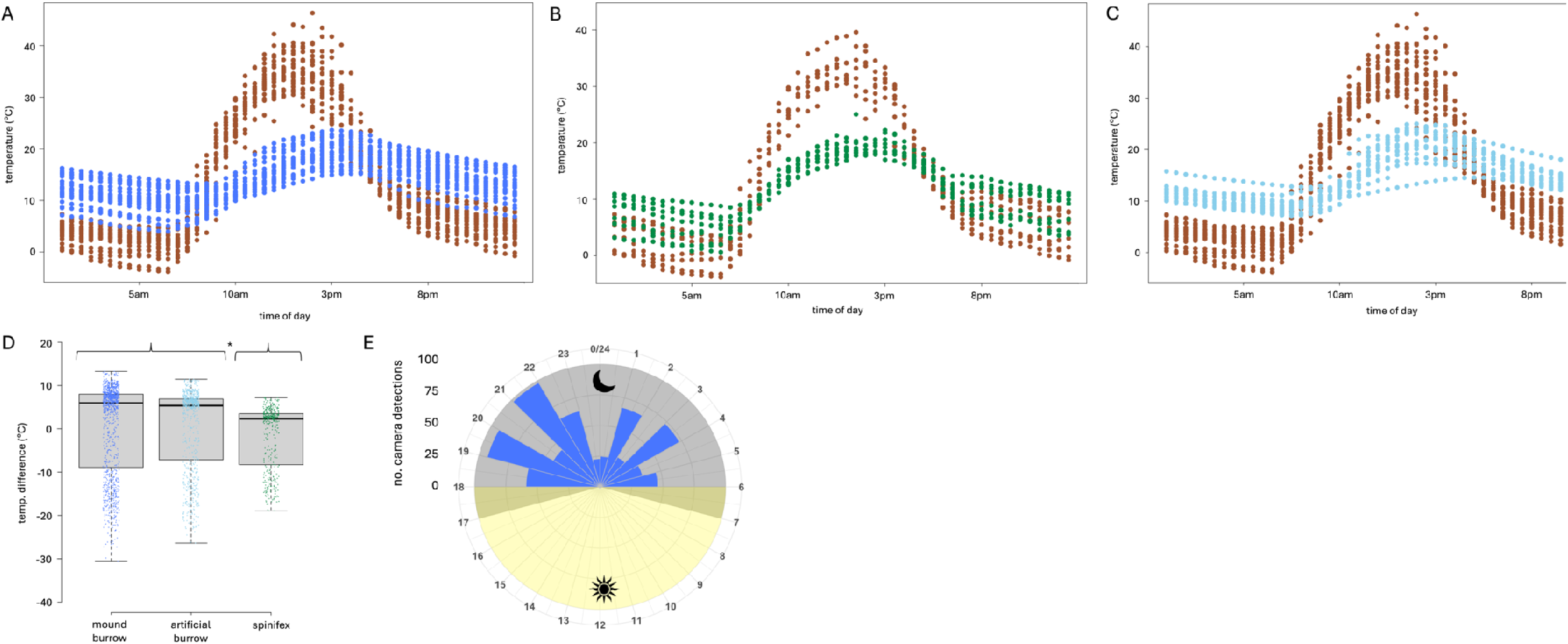
Winter ground temperatures (brown points) overlayed with mound burrow temperatures (A), spinifex shade temperatures (B) and experimental burrow temperatures (C). The difference between the ground temperature and each of the shelter types is displayed (D). The radar plot displays pebble mouse activity for four weeks overlapping the thermo-logging period (E).

**Figure 3.**
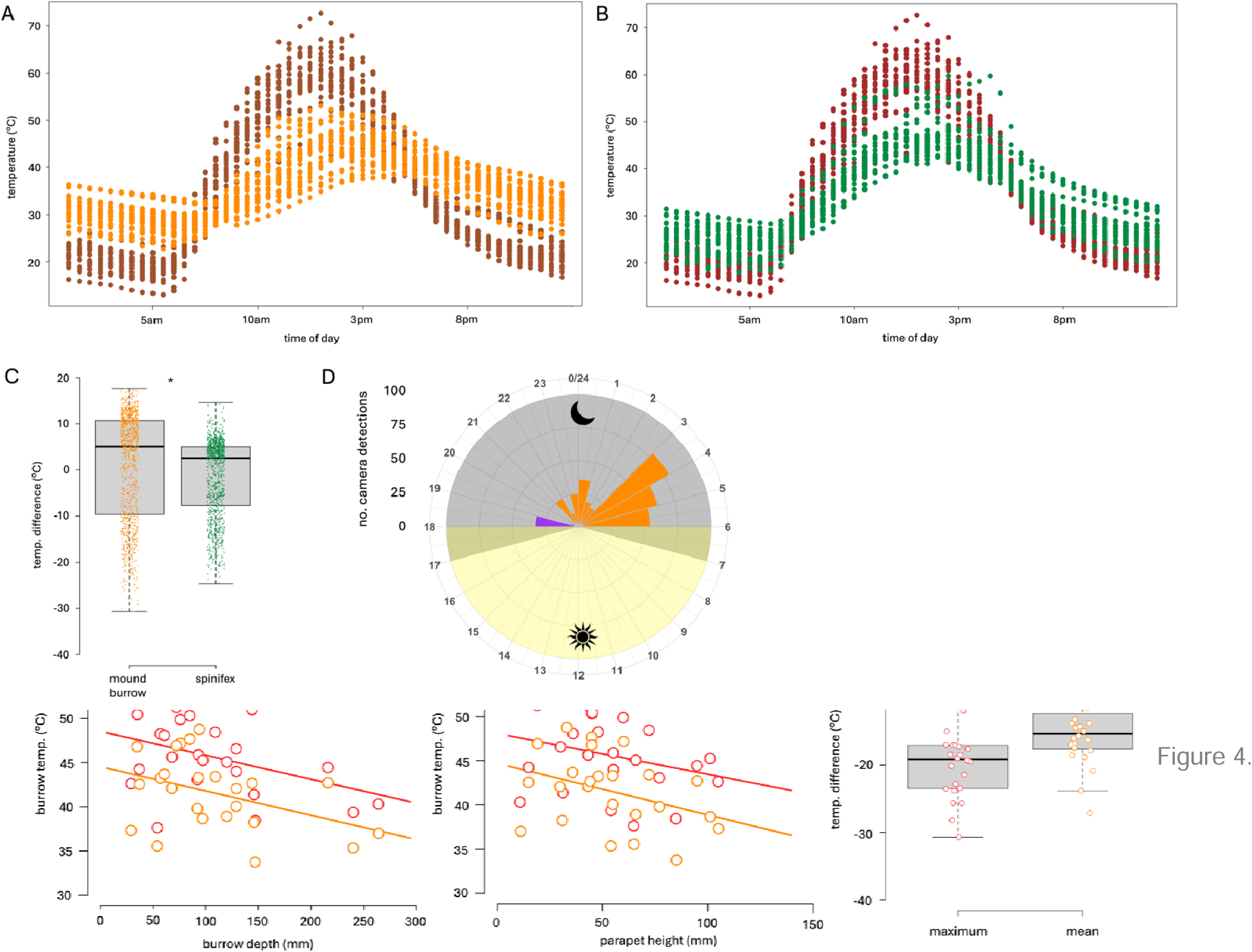
Spring ground temperatures (brown points) overlayed with mound burrow temperatures (A) and spinifex shade temperatures (B). The difference between the ground temperature and each of the shelter types is displayed (C). The radar plot displays pebble mouse activity for four weeks overlapping the thermo-logging period (D).

A total of 655 detections (images) were recorded in winter and 335 detections (images) in spring. The activity of pebble mice differed at the different times of the year with a predominance of activity before 22:00 in winter (Fig. 2E), but not spring (Fig. 3D). Thus, in winter most activity was recorded when the overnight ground temperatures were the warmest (19:00 to 22:00 hours at 10°C) and a large reduction in activity when the temperatures were the coldest (05:00 hours at below freezing to 5°C) (Fig. 2A, E). In spring, mouse activity did not overlap with observed Stimpson python activity (18:00 to 20:00 hours) and was mostly restricted to the last three hours of the night when it was coolest (03:00 to 06:00 hours at 25°C) (Fig. 3A, D). In the winter camera trap images, we observed that mice were often positioned on top of the mound, giving the impression that they were moving around the mound surface. In the spring images mice were typically captured either entering or exiting a burrow with little evidence of movement occurring on top of the mound surface.

### (b) Temperature and mound characteristics

The descriptive statistics for the mound characteristics are presented in Table S1, which show comparable measures for mounds logged in winter and spring. Across the 48 mounds, there were fewer northern facing burrows (i.e., 330° to 29°) compared to counts within the other categories (total no. burrows: 330° to 29° = 4; 30° to 89° = 10; 90° to 149° = 10; 150° to 209° = 8; 210° to 269° = 8; 270° to 329° = 8), though the observed values did not differ significantly from the expected distribution in this small dataset (df = 5, X^2^ = 3.000, p = 0.700).

We were interested in whether the magnitude of temperature buffering differed between the three shelter types (mound burrow, experimental burrow, spinifex shade). The analysis for winter revealed that there were differences among the shelter types (whole model: df = 2, X^2^ = 105.72, p < 0.001) with a post-hoc analysis providing further information that both the mound and experimental burrows had comparable buffering capacities (mound burrow vs. experimental burrow: *z*-value = - 1.325, p = 0.185), which was greater than the spinifex shade (mound burrow vs. spinifex shade: *z*-value = -10.242, p < 0.001; experimental burrow vs. spinifex shade: *z*-value = -7.702, p < 0.001) (Fig. 2D). Similarly, in spring, the mound burrows had a greater buffering capacity than the spinifex shade (df = 1, X^2^ = 127.55, p < 0.001) (Fig. 3C).

After accounting for variation due to outside ground temperature, our analyses revealed that burrow depth and parapet height were general predictors of variation in burrow temperature (Table 1). For burrow depth, this was evident across all hours in both winter and spring, but evident only in the cold hours in winter and only the hot hours in spring (Table 1). For parapet height, this was evident for the cold hours in winter and across all hours and the hot hours in spring (Table 1). Visualisation of the data revealed that deeper burrows and taller parapets were associated with keeping the burrows warmer in winter during the cold hours (Fig. 4A, B, C) and cooler in spring during the hot hours (Fig. 4D, E, F). Burrow angle was also important for keeping burrows warm during winter (all hours and cold hours) (Table 2, electronic supplementary material Fig. S4).

**Table 1.** Linear mixed models (LMMs) testing whether different mound characteristics accounted for variation in mound-burrow temperatures (after controlling for outside ground temperature). These analyses were performed on the entire dataset, which included “all hours” (i.e., 00:00 to 23:00), as well as in data subsets that included only the “cold hours” (i.e., 20:00 to 05:00) or the “hot hours” (i.e., 10:00 to 15:00).

|  | Chisq | df | p-value |
| --- | --- | --- | --- |
| (a) Winter |  |  |  |
| (i) Mean burrow temperature: all hours |  |  |  |
| Ground temperature | 19.628 | 1 | **<0.001** |
| Burrow depth | 17.007 | 1 | **<0.001** |
| Parapet height | <0.001 | 1 | 0.996 |
| Angle | 12.836 | 1 | **<0.001** |
| Aspect | 0.003 | 1 | 0.987 |
| (ii) Mean burrow temperature: cold |  |  |  |
| Ground temperature | 22.373 | 1 | **<0.001** |
| Burrow depth | 42.324 | 1 | **<0.001** |
| Parapet height | 4.608 | 1 | **0.032** |
| Angle | 25.065 | 1 | **<0.001** |
| Aspect | 0.029 | 1 | 0.864 |
| (ii) Mean burrow temperature: hot hours |  |  |  |
| Ground temperature | 0.647 | 1 | 0.421 |
| Burrow depth | 0.159 | 1 | 0.690 |
| Parapet height | 0.426 | 1 | 0.514 |
| Angle | 0.735 | 1 | 0.391 |
| Aspect | 0.629 | 1 | 0.428 |
| (b) Spring |  |  |  |
| (i) Mean burrow temperature: all hours |  |  |  |
| Ground temperature | 1.310 | 1 | 0.252 |
| Burrow depth | 9.351 | 1 | **0.002** |
| Parapet height | 7.861 | 1 | **0.005** |
| Angle | 3.446 | 1 | 0.063 |
| Aspect | 0.002 | 1 | 0.963 |
| (ii) Mean burrow temperature: cold |  |  |  |
| Ground temperature | 0.340 | 1 | 0.560 |
| Burrow depth | 0.011 | 1 | 0.918 |
| Parapet height | 0.158 | 1 | 0.691 |
| Angle | 0.571 | 1 | 0.450 |
| Aspect | <0.001 | 1 | 0.986 |
| (ii) Mean burrow temperature: hot hours |  |  |  |
| Ground temperature | 2.061 | 1 | 0.151 |
| Burrow depth | 7.686 | 1 | **0.006** |
| Parapet height | 4.678 | 1 | **0.031** |
| Angle | 1.319 | 1 | 0.251 |
| Aspect | 0.013 | 1 | 0.911 |

**Figure 4.** The relationships between burrow depth and parapet height and mound burrow temperatures during the cold hours (8pm to 5am) in winter (A, B; light blue: minimum temps, dark blue: mean temps) and the hot hours (10am to 3pm) in spring (D, E; orange: mean temps, red: maximum temps). The maximum and mean temperature difference between the mound burrow temperatures and ground temperatures are displayed (C, F).

### (c) Past and current soil temperatures

Our analyses revealed that the historical and the thermo-logged data were different in both months and across all hours, as well as in the subsets of only the cold and hot hours (Table S2, Fig. 5). The plotted temperatures and the mean differences show that thermo-logged temperatures were typically cooler in winter and warmer in spring except for the period of 14:00 to 16:00 in winter, where the historical and thermo-logged ground temperatures were comparable, and that the hot hours in spring where thermo-logged ground temperatures were cooler than historical ground temperatures (Fig. 5).

**Figure 5.**
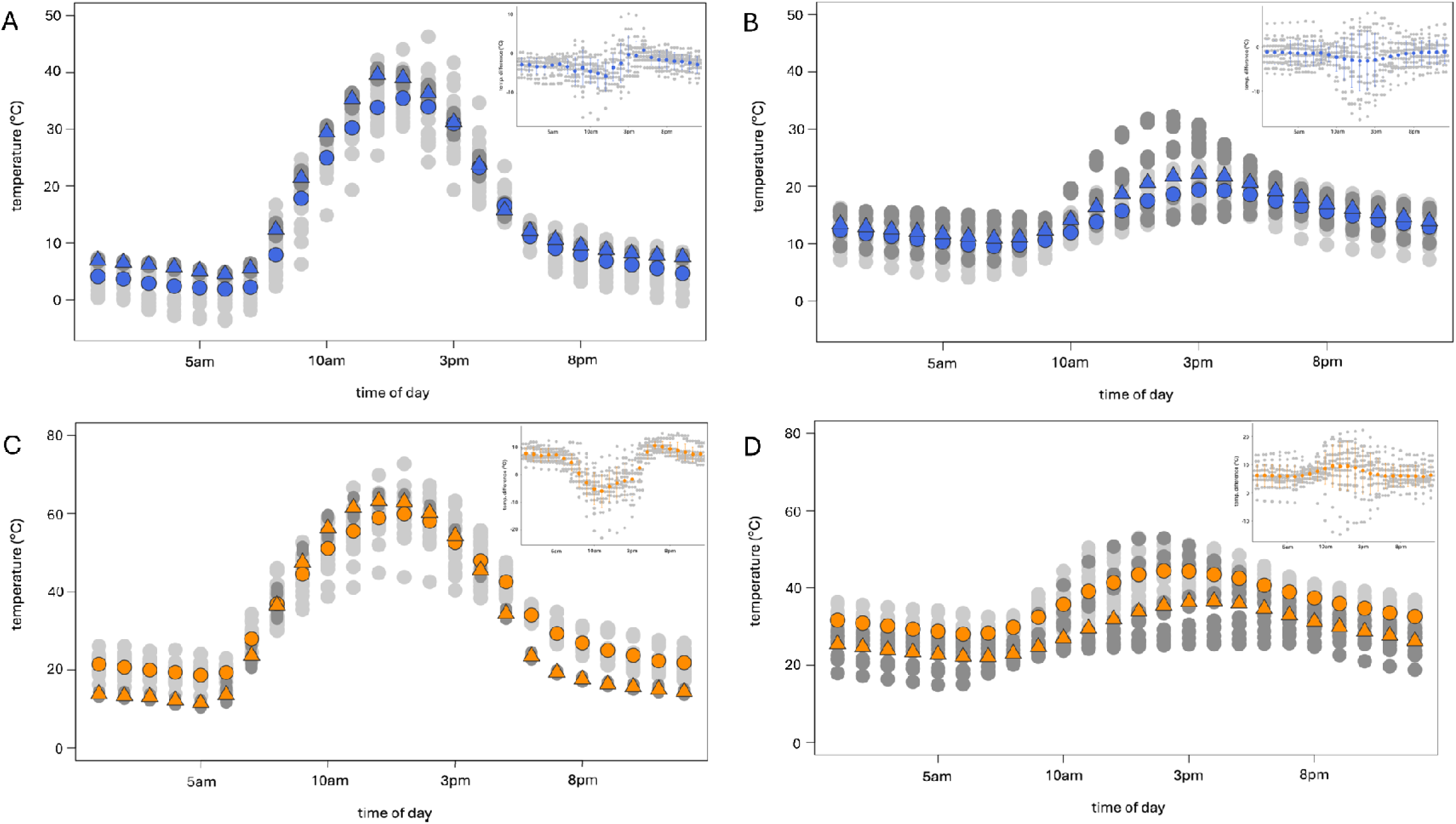
Winter (A, B) and spring (C, D) ground and mound burrow for data recorded in this study (light grey) and historical data (1960-1990) generated by NicheMapR (dark grey). The mean values are displayed (circles: thermo-logged data; triangles: historical data). The inserts show the temperature difference (light grey) with the mean ± s.e. displayed (coloured points and error bars).

### (d) Past, current and future thermoregulation

The thermoregulation models revealed that pebble mice are unable to survive at ground level (without shelter) during the peak heat of the day in any of the three months; the exceptions being in small individuals that (i) could survive at ground level under past winter conditions and (ii) did not quite reach the fatal limit under current winter conditions (Fig. 6). Overall, the winter models – past, present and future – show that if an individual is underground (≥10cm) they will remain normothermic (i.e., maintain a normal body temperature) across the entire day (Fig. 6B). In spring, the present condition models show a similar pattern to past conditions, with emergence to the surface after sunset allowing survival. However, it is evident that this timeline is delayed under current conditions (i.e., 17:00 vs. 18:00/19:00 hours) (Fig. 6 A, B) and under future conditions this timeline is delayed further (i.e., 20:00 hours) (Fig. 6C). The models show that in the past and currently, individuals ≥20cm underground will survive the peak heat of a spring day (Fig. 6 A, B). In the future, smaller individuals will be able to persist at this depth but larger individuals will need to be ≥50cm below surface level to remain normothermic during the hot hours (Fig. 6C). The summer models show that in the past and currently, individuals can survive the hottest month of the year at a depth of 50cm underground (Fig. 6A). However, according to the model, under current conditions larger individuals are unable to maintain a stable core body temperature at this depth and are required to persist at ≥100cm (Fig. 6A). In the future, large individuals will be unable to remain normothermic even at 100cm below ground level (Fig. 6A).

**Figure 6.**
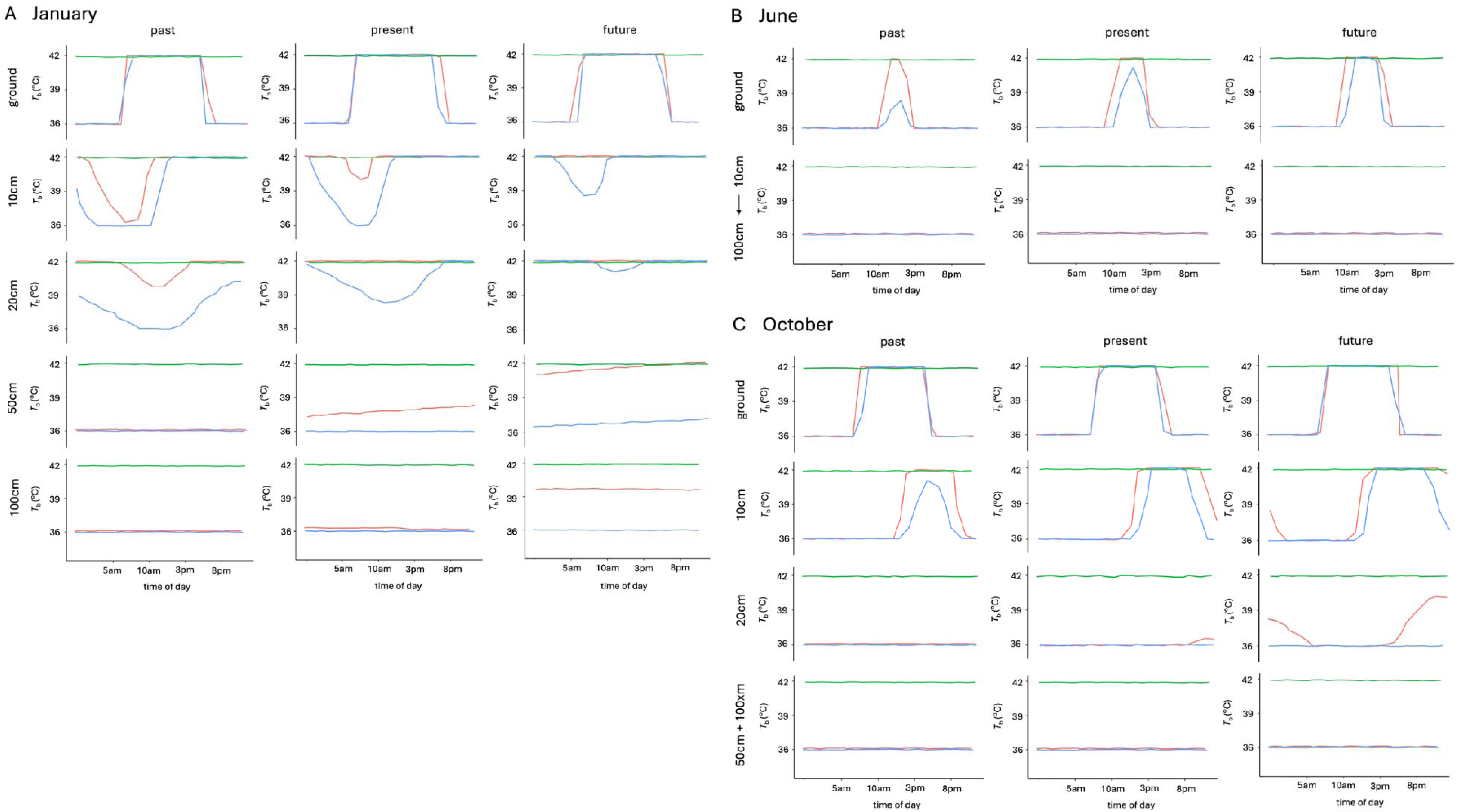
Core body temperatures (*T*_b_) modelled by NicheMapR under historical conditions (1960-1990), present conditions (historical+1.5°C) and predicted future conditions (historical+3°C) for summer (A January), winter (B June) and spring (C October). Coordinates of a mound-burrow system at approximately the midpoint of the study site (-22.61989°, 118.35638°) were applied. Values are provided for larger (18g; pink) and smaller (12.5g; blue) individuals. The green line represents the inability to thermoregulate (death).

The thermoregulation models generated with warming conditions of 4, 5 and 6°C showed little variation in profiles across the three temperatures (within months) and similar patterns to conditions modelled at +3°C warming (i.e., compare Fig. 6 to electronic supplementary material Fig. S5) as described above.

## 4. Discussion

Burrowing underground provides numerous benefits to animals, particularly in extreme environments where temperature regulation, water conservation, and protection from predators are critical for survival. One of the primary advantages is thermal insulation, as subterranean habitats can buffer against extreme surface temperatures, reducing heat exposure during the day and retaining warmth at night [42]. This is especially true for small mammals, such as the western pebble mouse, that live in arid or semi-arid environments where surface temperatures reach lethal levels. Here, we demonstrate that compared to having no shelter (i.e., outside ground temperature), or to sheltering under a bush, pebble mouse burrows were significantly warmer in winter and cooler in spring. We thus show that the pebble mound-burrow system, as an extended phenotypic trait, is a critical resource that creates a relatively stable microclimate and protects individuals from fluctuations in extreme surface conditions throughout the year.

Our investigation revealed that a unique mound characterisitc, the height of the pebble parapet around the burrow entrance, was associated with burrow temperatures – the taller the parapet the warmer the burrow in winter and the cooler the burrow in spring. Taller savanna termite mounds maintain higher constant internal temperatures than shorter mounds [43], and studies on the ventilation properties of burrows bearing a structure surrounding the entrance, such as a mound or chimney, have shown that height of these structures contributes to airflow into the burrow [44, 45]. Thus, taller pebble parapets may be more efficient at forcing moving air up and over the burrow entrance, rather than flowing into the burrow, resulting in warmer burrows in winter and cooler burrowers in spring. In winter, however, the experimental burrows (i.e., that lacked a pebble parapet) had comparable thermal profiles to the mound-burrows. This suggests that the magnitude of warming was not sufficient to impact the overall thermoregulatory properties of the mound-burrow system. Indeed, the excavation of experimental burrows in winter revealed that tunnelling into the earth, irrespective of whether the burrow was engineered by an animal or amassed with a pebble mound, was sufficient for offsetting early morning sub-zero temperatures. This perhaps isn’t too surprising considering that the Pilbara earth has heat retaining properties by being dark-red brown in colour and heavily concentrated with fragments of iron ore.

As observed in burrows engineered by other mammals [16–19], we discovered that depth underground was associated with the degree to which burrows were warm in winter and cool in the warmer months. While the maximum depth of the mound-burrow systems is yet to be scientifically investigated, an anecdotal report suggests that these structures, which have multiple tunnels and chambers, may extend to 60cm below the surface [34]. Indeed, burrows may turn and bend, and it is possible that our thermo-loggers did not reach the maximum depth. The relationships that we observed indicate that depths beyond 20–30cm (i.e., the maximum logged here) would have further warming (winter) and cooling (spring) properties. We suggest that the mice may persist at different depths underground depending on the prevailing outside conditions and how these influence the temperature within the burrow.

Burrows can also aid in moisture retention [12], which is important for arid-dwelling animals that are susceptible to desiccation and dehydration. The optimal range of humidities for dehydration avoidance in laboratory mice has been shown to be between 5–15g/m³ (at 20-25°C; [46]), which was similar to the range of subterranean humidities (within both mound and experimental burrows) that we observed in winter (peaking at 20g/m³; see electronic supplementary material). This provides evidence of buffering from the dry conditions of the cold winter air, which would be especially critical for breeding females and their need to maximise water retainment to support lactation and elevated heat production (as seen in the desert-dwelling, spiny mouse (*Acomys cahirinus*); [47]). The range of burrow humidities was wider in spring, falling as low as 0g/m³ but with the upper limit still at ∼15g/m³ (see electronic supplementary material). Notably, as external humidity increased during the hot hours, humidity levels within burrows remained relatively low and stable, which could offer several benefits. Primarily, exposure to lower and consistent humidities whilst taking refuge in the burrow would aid in thermoregulation (dissipation of body heat through evaporation) and reduce the risk of dehydration by offsetting water loss. Further, humidity levels maintained at an optimal level can contribute to the structural integrity of the burrow, ensuring that they remain safe and functional for shelter [48].

Due to the dry and powdery nature of the soil, we were unable to create experimental burrows in spring; an issue that western pebble mice may also encounter. Indeed, as suggested by variation in the intensity of activity centred on the mound in winter and spring, effective construction may be restricted to specific times of the year. While the energetic cost of construction has not been quantified, the size of the pebbles that the mice move (up to 6g) is indicative a substantial investment. Under certain conditions, individuals may live in social groups, which facilitates cooperation in burrow excavation and mound formation among group members [31]. Our work to date, however, shows that mounds are usually inhabited by a single, large female and it is these individuals that we observe moving pebbles (R, pers. obs.). This sex difference in mound building behaviour could at least partially explain why females are larger than males in this species. Certainly, the expansive size (up to 7m^2^) and suggested complexity of the mounds is likely due to the activity of multiple generations of mound inhabitants.

The thermologged mound-burrow data in spring often peaked at tempertures over 40°C, which is known to be close to the upper thermal tolerance limit of the arid-adapted cactus (*Peromyscus eremicus*) and canyon mouse (*Peromyscus crinitus*) (43–45°C) [49]. These *Peromyscus* species inhabit North American deserts where they are subjected to a *T*_a_ range that is comparable to the Pilbara region of Western Australia, and are a comparable size to pebble mice (16–21g) [49], which implies that pebble mice have similar heat tolerance limits. If this is the case, pebble mice would need to persist at cooler temperatures (deeper burrows) during the warmer months and especially during the peak of summer (January). Under current spring conditions, our modelling revealed that a depth of 20cm underground (i.e., similar to the maximum depths that we logged) ensures that individuals remain normothermic during the hot hours. The summer model for present day, however, indicates a “safe” depth of ∼50cm, supporting the notion that the mound-burrow system extends beyond the depths that we were able to monitor in this study. An assessment of the thermoregulatory performance of *Peromyscus* mice exposed to high *T*_a_ revealed an absence of effective cooling mechanisms, indicating that survival is almost entirely mediated by behaviour [49]. This may also be the case for western pebble mice with critical behavioural processes potentially including the construction and use of burrows or chambers at varying depths at different times of the year.

The Pilbara region is expected to increase by up to 3°C in the next 25 years [28, 29], which raises questions about how the fauna – many of which already persist at their thermoregulatory limits – will survive. Our soil microclimate hindcast modelling revealed that current ground and subterranean temperatures at our study site have changed relative to past conditions. We observed warmer ground and burrow temperatures in spring (except for moderately cooler ground temperatures from 10:00 to 13:00 hours) and cooler temperatures in winter. A comparison of the thermoregulation profiles for a past, present, and future winter are relatively consistent, whereby the high solar radiation and warming element of the substrate means that the burrows will continue to provide thermal protection from the cold nights. According to the model, future winter daytime temperatures should not be too impactful on the thermoregulation profiles of the mice. However, the predicted depth required to escape the heat of future summers (100cm) is striking. Even at this depth, large individuals, which are the breeding females, will be maintaining an elevated *T*_b_ of >39°C. Can western pebble mice burrow to depths exceeding half a metre, and potentially up to a metre, below the surface? Currently, we do not know, but answering this question appears to be critical to understanding whether this species will be able to persist under future climate change conditions.

Nocturnality is another behavioural trait that allows western pebble mice to persist in extreme Pilbara environment. Our hindcast modelling revealed a significant increase in nighttime ground temperatures (up to 5°C warmer). As we do not have activity data from the historic period, we are unable to draw conclusions about changes in activity that may have occurred in response to elevated nighttime temperatures. However, our current camera trapping data suggests that western pebble mouse activity may be driven, in part, by predator activity, and specifically the activity of Stimpson’s pythons. These nocturnal pythons are inactive in the coldest months of the year and most active when nighttime temperatures are between 26–30°C [50]. Thus, in winter, when there is no threat of pythons, activity occurs throughout the night. In spring, avoiding pythons also means that western pebble mice are active during the cooler part of the night (20–25°C), minimizing the metabolic demands of thermoregulation whilst foraging. Our forecast modelling suggests that future elevated temperatures of up to 5°C warmer are unlikely to impact the nocturnal thermoregulation profiles of the mice, at least in terms temperatures being “safe” for individuals to surface from the burrow. However, increasing temperatures may lead to shifts in the nocturnal behaviour of their predators, such as the Stimpson’s python, which could come at a huge cost – costs that may extend beyond the direct “consumptive” impacts of predation events and include nonlethal costs associated with the avoidance of predation risk (e.g., a reduction in foraging time) [51]. This reflects the importance of monitoring behavioural responses to climate change in predatory species when qualifying behavioural responses in prey species, such as rodents.

## 5. Conclusions

Our results indicate that the mound-burrow system is an essential modification of the western pebble mouse’s environment that buffers individuals from extreme temperatures, offering a stable, controlled environment that supports their thermoregulatory requirements. The thermoregulatory benefit must therefore contribute to offsetting the (likely large) cost of excavating the mound-burrow systems. Currently, however, the adaptive significance of the pebble mound (as an isolated entity) in the thermoregulatory properties of the system is not entirely clear. Our investigation revealed a significant correlation between height of the pebble parapet and burrow temperature, but this was not supported by the overall winter thermal profiles of burrows with (mound) and without (experimental) a pebble parapet. Indeed, the species’ reliance on burrowing into rocky areas of the eroding Pilbara hills may have little to do with thermoregulation, and alternate hypotheses accounting for the formation of the pebble mound should be investigated (e.g., refuge from flooding with the diversion of heavy cyclone-driven rainfall around burrow entrances).

If we align our findings with information on the thermal limits of other arid-adapted rodents, we can infer that western pebble mice likely rely heavily on behavioural processes to cope with searing hot and freezing cold conditions. However, we cannot discount other physiological processes that would aid in survival (e.g., daily torpor that has been observed in small desert rodents [52]). It would be useful to explore these potential physiological processes in the western pebble mouse, not only for the benefit of this species that has been poorly studied and faces multiple threats (i.e., anthropogenic disturbance, introduced predators, climate change), but to also gain a broader understanding of how small rodents, more generally, tolerate arid and semi-arid conditions.

Finally, our thermoregulation modelling under conditions of regional warming demonstrates the role that the extended phenotype can play in buffering the impact(s) of climate change. A critical consideration is whether individuals will be able to adjust their extended phenotype in an adaptive way, for example, in the case of the western pebble mouse, by being able to burrow deep enough underground to evade novel climatic extremes. Future work should aim to characterise this possibility, as well as explore potential costs, both direct and indirect, arising through changes in the behaviour or activity patterns of both target species and their predators.

## Ethics

This research followed the guidelines for the Code for the Care and Use of Animals for Scientific Purposes under Animal Ethics Committee approval (202 and permitted under a “Fauna taking (scientific or other purposes) licence” (F 0).

## Data accessibility

The data is provided as online supplementary material.

## Authors contributions

funding acquisition, project administration, conceptualization, investigation, data curation, formal analysis, methodology, validation, visualization, writing—original draft, writing—review and editing; funding acquisition, conceptualization, writing—review and editing.

## Acknowledgements

Karijini National Park is the traditional home of the Banjima, Kurrama and Innawonga Aboriginal people; this study was conducted in the southwest on Banjima Country. The authors acknowledge that the Banjima people are the traditional owners of the land on which this research was conducted, and we thank them for their support. We are extremely grateful to for assistance in the field, as well as and We thank for providing the humidity conversion equations. We also thank the Parks and Wildlife Karijini Rangers, especially and, for their assistance with our pebble mouse investigations.

